# Engineering assembloids to mimic graft-host skeletal muscle interaction

**DOI:** 10.1101/2024.11.20.622615

**Authors:** Lucia Rossi, Beatrice Auletta, Marco La Placa, Giada Cecconi, Pietro Chiolerio, Edoardo Maghin, Silvia Angiolillo, Eugenia Carraro, Onelia Gagliano, Cecilia Laterza, Nicola Elvassore, Martina Piccoli, Anna Urciuolo

## Abstract

Skeletal muscle (SkM) tissue engineering aims to generate in vitro three-dimensional (3D) products that can be implanted in patients to replace or repair damaged muscles. Having a humanized in vitro model able to mimic the interaction between the innervated recipient and the engineered SkMs at a functional level would greatly help in the evaluation of the graft potential.

Here we developed a 3D in vitro model that allowed to investigate the function, stability, and adaptability of the human NM system in response to an engineered SkM construct. To achieved this, we used decellularized SkMs (dSkM)-based constructs as engineered SkM and human neuromuscular organoids (NMOs) as the recipient-like NM system to create graft-host SkM assembloids.

We observed migration of myogenic cells and invasion of neural axons from the NMO to the engineered SkM construct in the assembloids, with the generation of functional neuromuscular junctions (NMJs). Finally, we showed that assembloids were able to regenerate following acute damage, with SkM regeneration and functional recovery.

Despite limited by the absence of immunocompetent cells and vasculature, our data showed that our assembloid represents a useful tool to evaluate in vitro the response of the human innervated SkM to a potential tissue-engineered SkM graft.

## Introduction

One of the main goals of tissue engineering is to generate in vitro organs and tissues that can be implanted in vivo to replace or repair diseased and damaged ones.[1] To date, terrific efforts have been made in this field with the generation of three-dimensional (3D) constructs capable of broadening knowledge on organ and tissue development and being used for translational applications.[2,3] While the precision and control over the generation of 3D in vitro constructs are being increasingly optimized thanks to progressively advanced technologies, there is a lack of information and investigation methods to predict the response of human cells to an engineered construct and subsequent interaction between the implant and host tissues. This is valid for any tissue type, including skeletal muscle (SkM).

In the last decade, several studies have been carried out for the in vivo replacement of large portions of SkM.[4–7] Current reconstructive options for extensive damage to SkM tissue or congenital SkM malformations include autologous muscle grafts and intramuscular injection of progenitor cells.[8] However, these therapeutic approaches are limited by many factors, such as the restricted availability of donor tissues (especially in pediatric patients), graft failure, donor site morbidity, and difficulties in regulating cell activity, which must be adequate and guided towards regeneration.[8] Moreover, SkM defects and damages are often accompanied by injury to the neuronal network directly connected to the muscle tissue, leading to atrophy and permanent functional disability of the denervated muscle, also after treatment.[9]

Several strategies have been developed in recent years to promote myogenesis and neuromuscular junction (NMJ) regeneration even after large volumes of muscle loss. Some tissue engineering approaches have included the use of customized scaffolds with controlled release of neurotrophic molecules,[10,11] also in combination with exercise,[4,10] to promote host-cell migration and/or invasion. It is known that scaffolds obtained through the decellularization of SkM (dSkM) possess intrinsic myogenic [5,12,13] and neurotrophic properties,[14] and when implanted in vivo as devices (i.e. without pre-seeding of muscular cells) can integrate with the host tissue in animal models and stimulate nerve sprouting, while simultaneously attracting myogenic progenitors inside the scaffold itself [12] with consequent SkM and NMJ regeneration.[13] Importantly, apart from evidence in animal models, it has been demonstrated that the implantation of decellularized scaffolds could promote a certain degree of muscle recovery in patients affected by volumetric muscle loss.[15] However, this and other studies [16,17] suggest that the use of decellularized scaffold as a medical device is not sufficient to guarantee full tissue recovery. A major challenge that hinders the complete functional integration of implanted scaffolds is the achievement of adequate and timely innervation by the host peripheral nervous system.[18] The use of dSkM pre-seeded with muscular cells (i.e. recellularized dSkMs, here defined as rSkMs) has been shown to be the most efficient solution to repair diaphragmatic hernia defects in animal models, as it can already provide a well-organized and functioning cytostructure [19] that can support the regeneration of the innervated SkM. However, the use of rSkM in patients affected by congenital or traumatic volumetric muscle loss conditions has never been reported, and no methods are currently available to predict the human cell response toward a recellularized construct in terms of myogenic regeneration, neuronal integration and functional NMJ formation. Indeed, at present, these humanized tissue-engineered constructs must be necessarily implanted in vivo in a xenogeneic environment to evaluate their integration properties, increasing the limiting aspects related to correctly predict human responses.[20,21]

Human neuromuscular organoids (NMOs) have been recently derived from both embryonic stem cells or human induced pluripotent stem cells (hiPSCs), enabling the production of in vitro models containing isogenic muscular and neuronal compartments arranged into spheroids.[22–25] The neuronal and muscular components in NMOs develop in parallel, self-compartmentalize, and interact to form functional NM networks with the formation of NMJs and consequent myofiber contraction.[22,24,26–28] Such NMOs have been therefore used to model and study in vitro the human NM system in health and disease. [22–25] We also recently derived tissue-engineered NMOs by the direct seeding of hiPSCs onto dSkMs, which offered a 3D extracellular environment able to promote the maturity of the NMOs and to mimic in vitro Duchenne Muscular Dystrophy myogenic phenotypes.[29] Moreover, Shin and colleagues have recently shown that SkM organoids derived from hiPSCs could be used to mimic and study in vitro myogenesis and regeneration of the human SkM upon acute damage with cardiotoxin.[24] Notably, the described SkM organoids also possessed a neuronal compartment.[24] Due to recent reports, the NMO technology is now considered a valuable tool to study and predict responses of the human NM system to damages and/or treatments.

Here we aim to study in vitro the human NM system response toward an engineered SkM construct that could be applicable in regenerative medicine. To do so, we combined NMO technology with a SkM tissue engineering approach, to develop a graft-host SkM assembloid model where the interaction of engineered constructs and the human NM system and their response to acute damage could be studied. Specifically, assembloids are in vitro models that combine two or more organoids, spheroids, or cultured cell types to recapitulate structural and functional properties of an organ or a system.[30] We used dSkM-based products as SkM tissue-engineered donor grafts and NMOs as recipient compartments to derive the engineered assembloids. We found that pre-seeding human muscle progenitors (hMPCs) within the dSkM (recellularized SkM, rSkM) promoted NMO host-cell invasion of the engineered constructs, when compared to scaffolds provided as device (dSkM - i.e. without pre-seeded hMPCs), with functional integration and NMJ formation. Furthermore, we subjected NMO-rSkM assembloids to acute damage via cardiotoxin treatment and we found that the assembloid was able to regenerate, rescuing the SkM function and partially the NMJs.

Implications of our research are significant and may contribute to advancements in tissue engineering and regenerative medicine of the human NM system, opening new perspectives for the translation of engineered SkM grafts to patients. Moreover, the possibility of faithfully analyzing the interaction between the human NM system and engineered SkM and vice versa represents an intriguing biological aspect that can increase our knowledge of the human NM system regeneration.

## Results

### Assembloids can be derived by combining dSkM scaffolds or rSkM tissue-engineered constructs and NMOs

With the aim to investigate the integration of NM system of human origin and engineered muscle constructs, we initially cultured NMOs derived from human induced pluripotent stem cells (hiPSCs)[29] with dSkM scaffolding biomaterials (NMO-dSkM assembloids) or with 7-day rSkM constructs [31] (NMO-rSkM assembloids; Fig. 1a). Along the entire study we alternatively used wild-type hiPSCs and hiPSCs constitutively expressing green fluorescent protein (GFP) under the control of 3-phosphoglycerate kinase promoter to derive NMOs (Supplementary Fig. 1). We used GFP-NMOs to easily follow the localization of cells derived from NMOs within our engineered SkM constructs, while wild-type NMOs were used for dedicated imagining analysis where the GFP signal could interfere - including Fluo-4 live imaging and immunofluorescence analysis. Upon selection and sorting of GFP-hiPSCs when necessary (Supplementary Fig. 1a, b), cells were characterized for pluripotency-related markers (Supplementary Fig. 1c-e) and used to produce self-assembled spheroidal NMOs (Supplementary Fig. 2). NMOs were derived using a small molecule-based differentiation protocol in the presence of Matrigel® droplets until day 22 (D22) from the beginning of the differentiation period [29] (Supplementary Fig. 2a). As first, we confirmed that GFP expression was still present at this stage of NMO differentiation (Supplementary Fig. 2b). We found that D22 NMOs showed muscular and neuronal compartmentalization, with a neuronal network that sprouted toward the myogenic components formed by elongated desmin (DES)-expressing cells (Supplementary Fig. 2c,d). As shown by immunofluorescence analysis, D22 NMOs also possess both paired box protein 7 (PAX7)-expressing stem cells and myogenic committed myogenin (MYOG)-positive progenitor cells, associated to myofibers that express embryonic myosin heavy chain (eMHC) in contact with laminin (LAM) deposited in the extracellular matrix (Supplementary Fig. 2e-h).

**Figure 1.**
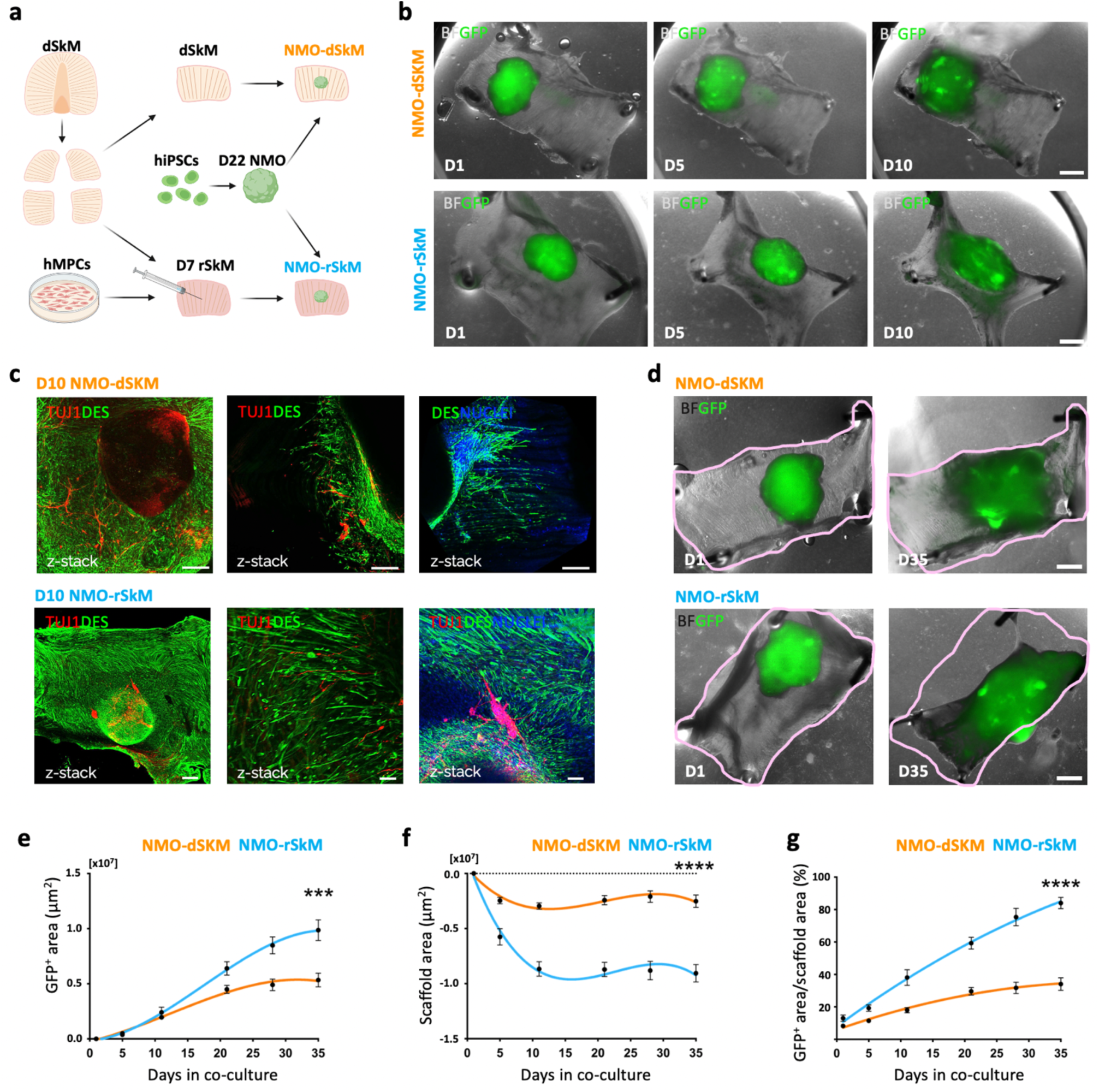
NMOs integrate within dSkMs and rSkMs to form graft-host SkM assembloids. **a**. Schematic illustration showing the strategy used for co-culturing NMOs and dSkMs (NMO-dSkM, orange) or rSkMs (NMO-rSkM, light blue). NMOs were derived from hiPSCs upon 22 days (D22 NMO) of NM differentiation. rSkMs were generated upon hMPCs injection within dSkM 7 days (D7 rSkM) before NMOs seeding. Acronyms list: dSkM: decellularized skeletal muscle; hMPCs: human muscle progenitor cells; hiPSCs: human induced pluripotent stem cells; D22 NMO: day 22 neuromuscular organoids; D7 rSkM: day 7 recellularized skeletal muscle construct; NMO-dSkM: construct obtained from NMO combined with dSkM; NMO-rSKM: construct obtained from NMO combined with rSkM. **b**. Representative brightfield (BF) and fluorescence stereomicroscope images showing the progression of GFP-NMOs at day 1 (D1), day 5 (D5) and day 10 (D10) after seeding onto dSkM (NMO-dSkM, upper panels) or onto rSkM (NMO-rSkM, lower panels). Scale bars, 1 mm. **c**. Upper left and middle panels, representative Z-stack confocal immunofluorescence images of whole mount NMO-dSkM after 10 days of co-culture (D10 NMO-dSkM) stained for DESMIN (green) and TUJ1 (red). Upper right panel, representative Z-stack confocal immunofluorescence image of whole mount NMO-dSkM after 10 days of co-culture (D10 NMO-dSkM) stained for DESMIN-positive cells (green) that invade the dSkM. Nuclei are counterstained with Hoechst (blue). Scale bars, 200 μm. Lower panel, representative Z-stack confocal immunofluorescence images of whole mount NMO-rSkM after 10 days of co-culture (D10 NMO-rSkM) stained for DESMIN (green) and TUJ1 (red). Right panel, nuclei are counterstained with Hoechst (blue). Scale bars, 500 μm (left), 50 μm (center), 100 μm (right). **d.** Representative brightfield (BF) and fluorescence stereomicroscope images showing the expansion of GFP-NMOs after 1 day (D1) and 35 days (D35) from seeding onto dSkM (NMO-dSKM) or rSkM (NMO-rSKM). The pink line is used to track the morphological changes of the overall construct shape at the two co-culture time points. Scale bar, 1 mm. **e.** NMO area variation (µm^2^) over culture time (days), expressed as difference between GFP^+^ area at day n and GFP^+^ area at day 1 of co-culture. Data are shown as mean ± SEM of n>9 independent replicates. Statistical significance determined using Mann-Whitney U test; ***P=0.0008 on day 35 (0.99×10^7^±0.09×10^7^ μm^2^ vs 0.53×10^7^±0.06×10^7^ μm^2^). **f.** Scaffold area variation (µm^2^) over culture time (days), expressed as difference between scaffold area at day n and scaffold area at day 1 of co-culture. Data are shown as mean ± SEM of n>9 independent replicates. Statistical significance determined using Mann-Whitney U test; ****P<0.0001 on day 35 (-0.91×10^7^±0.07×10^7^ μm^2^ vs -0.25×10^7^±0.06×10^7^ μm^2^). **g.** Ratio between NMO area and scaffold area, expressed in percentage. Data are shown as mean ± SEM of n>9 independent replicates. Statistical significance determined using Mann-Whitney U test; ****P<0.0001 on day 35 (84.02±3.52% vs 34.04±3.92%).

To produce tissue-engineered human SkM constructs, we used an already well characterized decellularization protocol to derive dSkM from the murine diaphragm,[32,33] which was used per se (dSkM) or recellularized (rSkM) with hMPCs isolated from healthy muscle biopsies.[34] We already demonstrated that dSKM sustains hMPC engraftment, proliferation and differentiation, generating 3D rSkM tissue-engineered constructs suitable for in vivo implantation.[19] Nevertheless, before being used to generate rSkM, we expanded and characterized hMPCs in culture (Supplementary Fig. 3a) confirming that they possess high cell rates of CD56 (94.18±2.22%) and MYOD (99.22±0.17%) expressing cells and low rates of MYOG (3.86±0.61%) positive cells (Supplementary Fig. 3b-d). In agreement with previous studies,[31,34] a small proportion of TE7^+^ (0.93±0.15%) fibroblasts were also identified in the primary hMPC culture (Supplementary Fig. 3e). Finally, to confirm the myogenic ability of hMPCs, we evaluated and quantified the proliferation rate and the ability to form myotubes in vitro (Supplementary Fig. 3f-g).

To understand the feasibility of the NMO/SkM-engineered assembloids, we first performed short-term experiments by seeding D22 GFP-NMOs onto dSkM or rSkM, and by culturing the assembloids for 10 days (Fig. 1b). In both cases (NMO-dSkM and NMO-rSkM assembloids), cells derived from NMOs integrated within the tissue-engineered products, as shown by the sprouting of both neural and muscular components observed in whole mount samples immunostained for TUJ1^+^ neuronal projection and DES^+^ myogenic cells, respectively (Fig. 1c). During the time of culture and at longer time points (35 days), we observed invasion of GFP-expressing cells within the scaffolds and a differential gross appearance of NMO-dSkM and NMO-rSkM assembloids (Fig. 1b,d and Supplementary Fig. 4a,b). In particular, we found that both tissue-engineered SkM and GFP-NMO shapes were evidently remodeled in NMO-rSkM after 35 days of co-culture when compared to NMO-dSkM assembloids at equivalent time points (Fig. 1d). To quantify such observation, we monitored NMO-rSkM and NMO-dSkM cultures, and we measured over time (from day 0 to day 35) the area occupied by GFP^+^ cells (Fig. 1e), the scaffold area variation (Fig. 1f) and their ratio (Fig. 1g). We confirmed that NMOs showed improved ability to spread, remodel and invade the scaffold when in contact with hMPC pre-seeded constructs - i.e. NMO-rSkM assembloids (Fig. 1e-g). Interestingly, we also found that hMPCs had per se the ability to remodel the dSkM during the time of culture - i.e. rSkM samples (Supplementary Fig. 4c,d). Altogether these data demonstrate that both dSkM and rSkM can be invaded by human NM system derived from the NMO in vitro, generating graft-host SkM assembloids.

### Pre-seeding of hMPCs into dSkMs promotes myogenesis and muscle functionality in NMO-rSkM assembloids

It remains a matter of discussion in the field whether the pre-seeding of tissue-engineered constructs with cells could promote the integration of the graft with recipient cells. Toward this aim, we compared NMO-dSkM and NMO-rSkM assembloids after 35 days of culture to understand whether the pre-seeding of hMPCs into dSkM could promote the integration of NMOs and overall myogenesis and muscle functionality of our in vitro assembloid model. As controls, rSkMs were included in the analysis to monitor sample behavior in the absence of NMO contribution. First, we performed morphometric analysis to evaluate the presence, localization and organization of differentiated myogenic cells within the samples. We confirmed the invasion of the scaffolds by seeded cells in all samples, as shown by nuclei and laminin immunofluorescence analysis (Supplementary Fig. 4e). Despite all samples showing elongated DES^+^ myogenic cells, we found that cells were extensively localized within the NMO-rSkM assembloids, identified as discrete bundles into NMO-dSkM assembloids, and as fewer cells within the rSkM samples (Fig. 2a,b and Supplementary Fig. 4f). Accordingly, with the expected developmental stages of the myotubes (i.e. from adult hMPCs or fetal/neonatal NMOs), we also found that rSkMs displayed statistically significant thicker myotubes (15.53±0.63 µm) when compared to NMO-dSkM and NMO-rSkM assembloids, with the presence of many but thin myotubes in the former (7.71±0.21 µm), and many and significantly thicker myotubes in the latter (10.32±0.26 µm; Fig. 2a,c).

**Figure 2.**
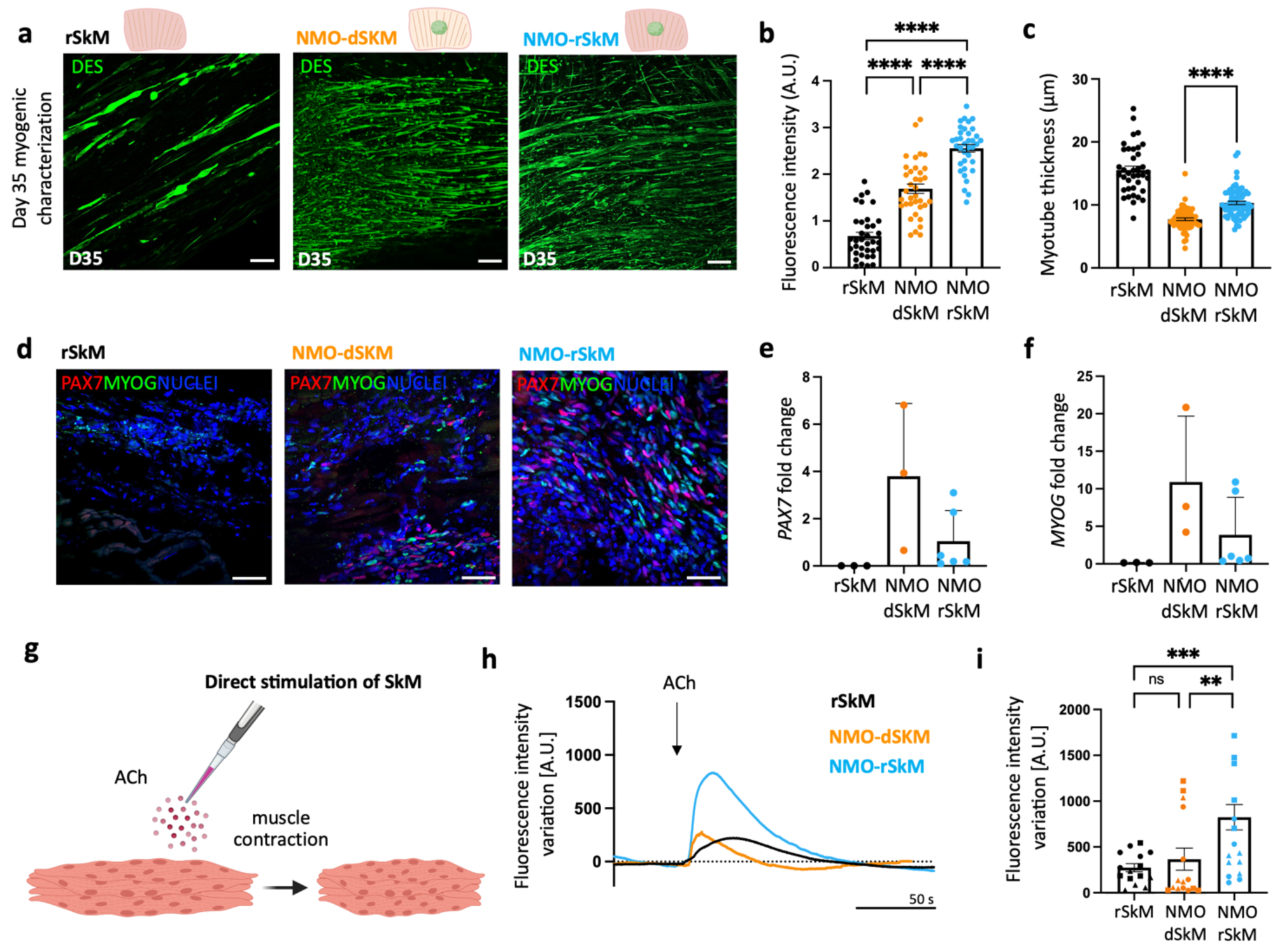
Myogenesis and muscle functionality are improved in NMO-rSkM assembloids. **a**. Representative Z-stack confocal immunofluorescence images of whole mount rSkM (left panel), NMO-dSkM (center panel) or NMO-rSkM (right panel) after 35 days (D35) of co-culture, stained for DESMIN (green). Scale bars, 100 μm. **b**. Quantification of corrected total fluorescence intensity normalized for background signal performed on DESMIN immunofluorescence staining. Data are shown as mean ± SEM of 36 ROIs per condition (3 images per condition, for each image 12 ROIs were selected and quantified). Statistical significance was determined using Mann-Whitney U test; ****P<0.0001. **c**. Quantification of DESMIN^+^ myotube cross-section (thickness) performed on whole mount immunofluorescence images of rSkM, NMO-dSkM and NMO-rSkM after 35 days (D35) of culture. Data are shown as mean ± SEM of >100 myotube cross sections, measured on 3 independent samples per condition; statistical significance determined using Mann-Whitney U test; ****P<0.0001. **d**. Representative confocal immunofluorescence image of rSkM, NMO-dSkM and NMO-rSkM cross-sections, stained for PAX7 (red) and MYOG (green). Nuclei are counterstained with Hoechst (blue). Scale bar, 50 µm. **e**. *PAX7* gene expression in rSkMs, NMO-dSkMs and NMO-rSkMs. Data are normalized to housekeeping *B2-microglobulin* gene expression and shown as fold change over NMO. Data are shown as mean ± s.d. of 3 independent biological replicates for rSkM and NMO-dSkM and of 6 independent biological replicates for NMO-rSkM. **f**. *MYOG* gene expression in rSkMs, NMO-dSkMs and NMO-rSkMs. Data are normalized to housekeeping *B2-microglobulin* gene expression and shown as fold change over NMO. Data are shown as mean ± s.d. of 3 independent biological replicates for rSkM and NMO-dSkM and of 6 independent biological replicates for NMO-rSkM. **g**. Schematic illustration showing the strategy used to assess muscle compartment functionality in response to exogenous ACh administration. **h**. Representative quantification of mean normalized fluorescence intensity variation registered during live imaging analysis of D35 rSkM, NMO-dSkMs and NMO-rSkMs, stimulated with ACh. Data are shown as mean of 9 ROIs, 3 ROIs were selected from each of 3 independent replicates. Dotted line corresponds to the baseline equal to 0. **i**. Contraction quantification of rSkM, NMO-dSkM or NMO-rSkM at D35 upon exogenous ACh administration expressed as maximum peak of fluorescence revealed via live imaging analysis. Quantifications were performed on 3 independent biological replicates, identified by different symbol shapes in the graph. Each independent sample was divided into 5 ROIs and each point represents a single ROI. Statistical significance was determined using Mann-Whitney U test; ns, not statistically significant, **P=0.0066, ***P=0.0001.

Apart from terminally differentiated myogenic cells, we also evaluated whether rSKMs, NMO-dSkMs and NMO-rSkMs could possess muscle stem cells (PAX7-expressing cells) and myogenic committed cells (expressing MYOG) after 35 days of culture. Immunofluorescence analysis showed that both NMO-dSkM and NMO-rSkM assembloids possess both stem and committed myogenic cells (Fig. 2d). The absence of PAX7 expressing cells and the small amount of MYOG expressing cells in rSkMs (Fig. 2d-f and Supplementary Fig. 4f-h) strongly suggested that NMOs are the major contributor to stem and committed myogenic cell populations revealed within NMO-dSkM and NMO-rSkM, maybe because of the long time point considered in this study (Fig. 2d-f, Supplementary Fig. 2e and Supplementary Fig. 4g,h).

Once confirmed that all samples possess a certain grade of myogenic commitment, and demonstrated that the pre-seeded scaffold triggers a wide myogenic differentiation and maturation, we evaluated the ability of these models to contract when stimulated with acetylcholine (ACh) by quantifying fluorescence intensity variation associated to muscle contraction during live imaging (Fig. 2g). Recellularized SkM samples, mainly composed of sparse mature myotubes with little interaction with scaffold extracellular matrix (Supplementary Fig. 4f), were able to respond to the exogenous ACh administration with a contraction intensity variation comparable to that of NMO-dSkM assembloids (273.65±43.79 A.U. vs 366.56±121.45 A.U.), which oppositely possess more but less mature myotubes (Fig. 2h-i, Supplementary Fig. 4i,j and Supplementary Video 1). On the contrary, NMO-rSkMs responded with significantly higher contraction to ACh administration when compared to the other samples (823.65±138.28 A.U.; Fig. 2h-i, Supplementary Fig. 4k and Supplementary Video 1). This was in line with characteristics of an increased proportion of the muscular compartment, when compared to rSkMs and NMO-dSkMs, and thicker myotubes when compared to NMO-dSkMs (Fig. 2b-d).

These results confirm the muscular integration of NMOs within tissue-engineered constructs to form graft-host SkM assembloids and demonstrate that the pre-seeding of hMPCs within dSkM promotes myogenesis and muscle functionality in NMO-rSkM.

### NMO-rSkM assembloids possess functional NMJs after 35 days of co-culture

Based on the above results, we focused our next studies on NMO-rSkM (Fig. 3a). The most limiting aspect in the in vivo application of muscle implants is the achievement of adequate and timely innervation by the host peripheral nervous system.[18] For this reason, we analyzed the presence of functional NMJs as an indication of a complete integration between NMOs and rSkM. From a general point of view, day 35 NMO-rSkM assembloids displayed strong and wide neurofilament (NF) positive axons which spread from the central body of the NMO to also distal parts of alpha-sarcomeric actin (αSA)-expressing cells (Fig. 3b). The NMO-rSkM assembloids appeared well integrated as shown by scanning electron microscopy (Supplementary Fig. 5a). Moreover, together with the expression of developing muscle markers (Supplementary Fig. 5b), we found the presence of supporting muscle cells such as TE7^+^ fibroblasts (Fig. 3c), deposition of basal lamina (Fig. 3d) and proteins essential for muscle contraction such as slow and fast MHC and sarcomeric TITIN (Fig. 3e,f). Importantly, we also identified juxtaposed pre- and post-synaptic elements of the NMJs in anatomically expected positions (Fig. 3g and Supplementary Fig. 5c-e). Moreover, in close contact between TUJ1^+^ neuronal protrusions and alpha-BTX^+^ signal, we discovered the presence of S100B^+^ cells (Fig. 3h and Supplementary Fig. 5e), strongly suggesting the presence of terminal Schwann cells.[22] All these data are also supported by the expression of genes that highlight the presence of early-stage MN (ISL1;[35]), and different isoforms of ACh receptors (ACHRγ and ACHRε; Fig. 3i-k), as indication of NMJ maturation.[36]

**Figure 3.**
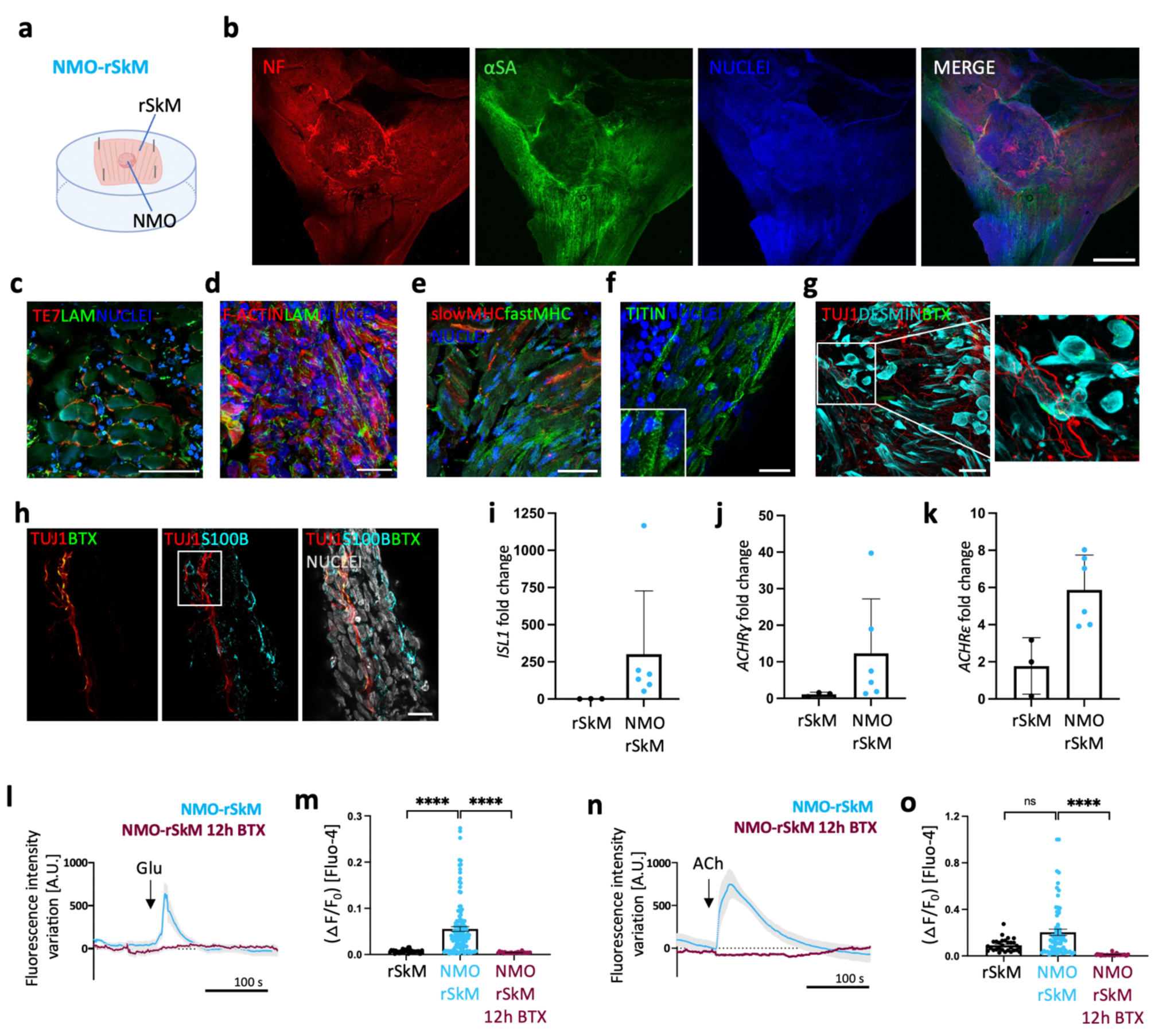
NMO-rSkM assembloids possess functional NMJs after 35 days of co-culture. **a.** Schematic illustration showing a representation of NMO-rSkM co-culture setup. **b**. Representative Z-stack confocal immunofluorescence images of whole mount NMO-rSkMs stained for NEUROFILAMENT (NF, red) and alpha-SARCOMERIC ACTININ (αSA, green). Nuclei are counterstained with Hoechst (blue). Scale bar, 1 mm. **c**. Representative Z-stack confocal immunofluorescence image of NMO-rSkM cross-sections stained for TE7 (red) and LAMININ (green). Nuclei are counterstained with Hoechst (blue). Scale bar, 50 µm. **d**. Representative Z-stack confocal immunofluorescence image of NMO-rSkM cross-sections stained for F-actin (red) and LAMININ (green). Nuclei are counterstained with Hoechst (blue). Scale bar, 20 µm. **e**. Representative Z-stack confocal immunofluorescence image of NMO-rSkM cross-sections stained for slow MYOSIN HEAVY CHAIN (MHC, red) and fast MHC (green). Nuclei are counterstained with Hoechst (blue). Scale bar, 50 µm. **f**. Representative Z-stack confocal immunofluorescence image of NMO-rSkM cross-sections stained for TITIN (green). Nuclei are counterstained with Hoechst (blue). Scale bar, 20 µm. Inset shows a higher magnification, evidencing sarcomeric organization of myofiber cytoskeleton. **g**. Representative Z-stack confocal immunofluorescence image of NMO-rSkM cross-sections stained for TUJ1 (red), DESMIN (cyan) and α-bungarotoxin^+^ (BTX) regions (green). Nuclei are counterstained with Hoechst (blue). Scale bar, 50 µm. **h**. Representative Z-stack confocal immunofluorescence image of NMO-rSkM cross-sections stained for TUJ1 (red), BTX (green) and S100β (cyan). Nuclei are counterstained with Hoechst (grey). Scale bar, 20 µm. **i**. *ISL1* gene expression in rSkMs and in NMO-rSkMs. Data are normalized to housekeeping *B2-microglobulin* gene expression and shown as fold change over rSkM. Data shown as mean ± s.d. of 3 independent biological replicates for rSkM and of 6 independent biological replicates for NMO-rSkM.. **j**. *ACHRɣ* gene expression in rSkMs and in NMO-rSkMs. Data are normalized to housekeeping *B2-microglobulin* gene expression and shown as fold change over rSkM. Data shown as mean ± s.d. of 3 independent biological replicates for rSkM and of 6 independent biological replicates for NMO-rSkM. **k**. *ACHRε* gene expression in rSkMs and in NMO-rSkMs. Data are normalized to housekeeping *B2-microglobulin* gene expression and shown as fold change over rSkM. Data shown as mean ± s.d. of 3 independent biological replicates for rSkM and of 6 independent biological replicates for NMO-rSkM. **l,n**. Representative quantification of mean normalized fluorescence intensity variation registered during live imaging analysis of NMO-rSkMs or NMO-rSkMs treated for 12 hours with BTX and stimulated with Glu (l) or ACh (n). Data are shown as mean ± SEM of 15 ROIs (5 ROIs per sample, from 3 independent biological replicates) for untreated NMO-rSkMs (light blue line). Data are shown as mean ± SEM of 10 ROIs (5 ROIs per sample, from 2 independent biological replicates) for NMO-rSkMs treated overnight with BTX (crimson line). Dotted lines correspond to the baseline equal to 0. **m,o**. Quantification of calcium peak amplitude (ΔF/F_0_) detected with Fluo-4 live imaging analysis of rSkM (n=2), NMO-rSkM (n=4) or NMO-rSkM after BTX treatment (n=2) after Glu (m) or ACh (o) administration. The fluorescence intensity peak (F) during stimulation was measured and normalized to the baseline fluorescence intensity registered before neurotransmitter stimulation (F_0_). Mann-Whitney U test; ns, not statistically significant; ****P<0.0001.

To test whether the identified NMJs were also functional, we stimulated NMO-rSkMs with glutamate (Glu) and evaluated the muscle contraction. In vivo, Glu stimulates the MN to release ACh at the NMJ, inducing muscle contraction. Only if the NMJ is properly formed, it is possible to induce muscle contraction through Glu administration. We found that neuronal-mediated stimulation of NMJs resulted in NMO-rSkM assembloids contraction and significant increase of calcium spikes in the muscular compartment, thus indicating the presence of functional NMJs (Fig. 3l,m, Supplementary Fig. 5f,g and Supplementary Video 2). Importantly, calcium transients were not revealed in rSkM samples, which do not possess a neuronal compartment, indicating the specificity of action of the Glu at the muscular level via the neuronal stimulation (Fig. 3m). As control of the physiological activity of the muscle compartment in our samples, we also supplemented ACh to reveal signals derived upon direct simulation of AChRs situated on myofibers. As expected, the addition of exogenous ACh was able to induce a positive response in terms of contraction and calcium transient in all the analyzed samples, including rSkMs (Fig. 3n,o and Supplementary Video 3). In agreement, 12h nicotinic AChR block with BTX led to the loss of NMO-rSkM contraction and calcium transients, both after stimulation with Glu or ACh, highlighting the physiological behavior of the newly formed NMJs (Fig. 3l-o, Supplementary Fig. 5g,h and Supplementary Videos 2-3).

Taken together, these data indicate that a functional NM integration between the NMO system and a tissue-engineered muscle construct was achieved in NMO-rSkM assembloids.

### NMO-rSkM assembloids possess muscle stem cells and model regeneration events upon acute damage

To define the integration between the NMO and the rSkM (i.e. NMO-rSkM) as definitely functional and potentially long-lasting, it is necessary to verify that the assembloid could respond to acute injury with regeneration. A relevant body of studies demonstrated the key role of PAX7 expressing cells for proper SkM regeneration.[37,38] After 35 days of culture, the NMO-rSkM assembloids presented both stem and committed myogenic cells expressing PAX7 or MYOG, respectively (Fig. 2d, Fig. 4a,b and Supplementary Fig 5b). Furthermore, these cells were found in well-defined anatomical positions, with PAX7^+^ cells also located beneath the basal lamina of the muscle fibers, which defines the anatomical position of satellite cells (Fig. 4c).[37,38] It should be noted that, at this culture time point, more than 90% of PAX7^+^ cells were not actively proliferating, and that only 6.28±1.66% co-expressed the proliferation marker Ki67 (Fig. 4d).

**Figure 4.**
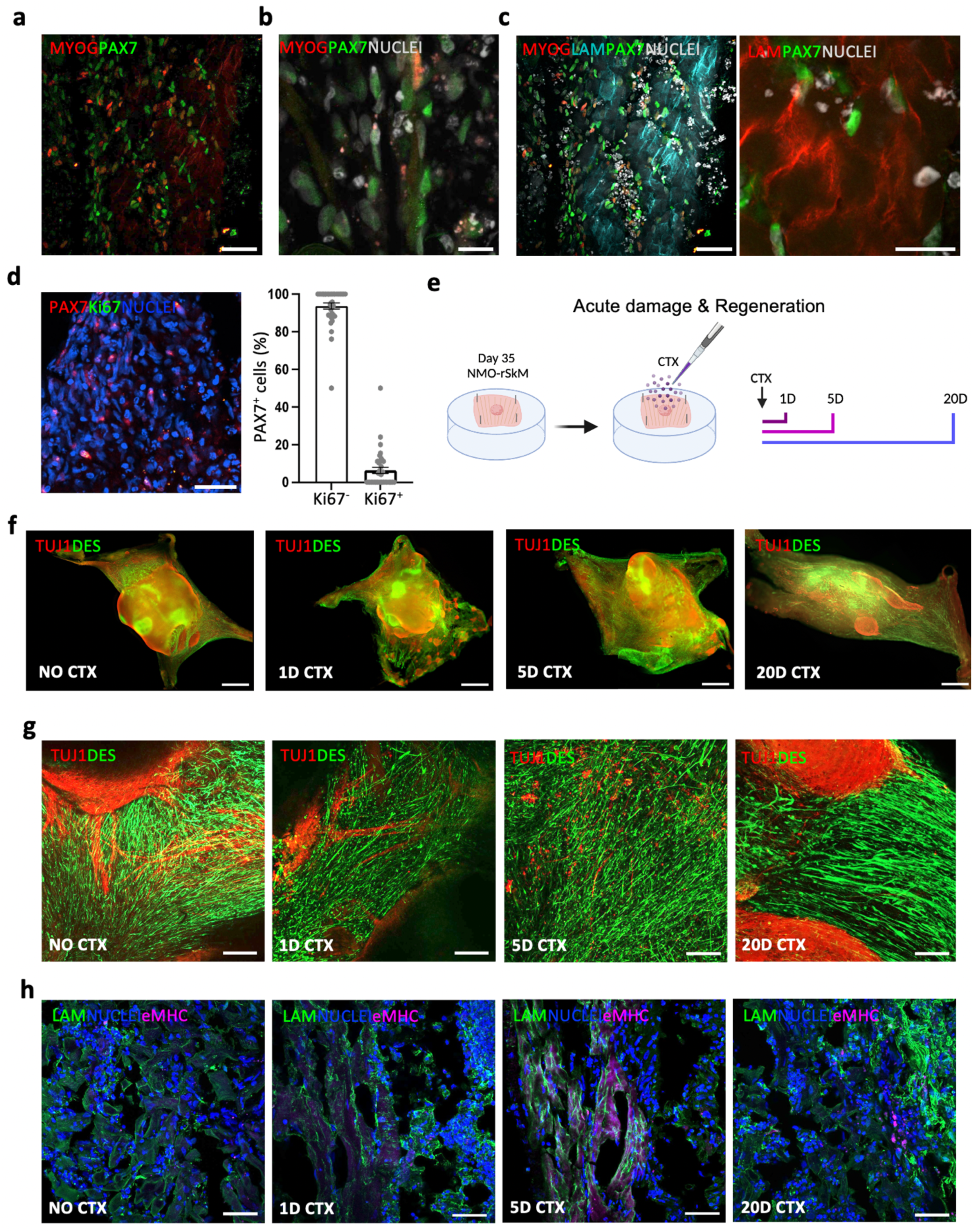
NMO-rSkM assembloids possess muscle stem cells and model degeneration and regeneration events upon acute damage. **a-b**. Representative Z-stack confocal immunofluorescence image NMO-rSkM cross-sections at day 35 of co-culture stained for MYOG (red) and PAX7 (green). Nuclei are counterstained with Hoechst (grey). Scale bars, 50 µm (a) and 20 µm (b). **c**. Left panel, representative Z-stack confocal immunofluorescence image of NMO-rSkM cross-sections at day 35 of co-culture stained for MYOG (red), LAMININ (cyan) and PAX7 (green). Nuclei are counterstained with Hoechst (grey). Scale bar, 50 µm. Right panel, representative Z-stack confocal immunofluorescence image of NMO-rSkM cross-sections at day 35 of co-culture stained for LAMININ (red) and PAX7 (green). Nuclei are counterstained with Hoechst (grey). Scale bar, 20 µm. **d**. Left panel, representative Z-stack confocal immunofluorescence image of NMO-rSkM cross-sections at day 35 of co-culture stained for PAX7 (red) and Ki67 (green). Nuclei are counterstained with Hoechst (blue). Scale bar, 50 µm. Right panel, quantification of PAX7^+^Ki67^+^ and PAX7^+^Ki67^-^ cells on total PAX7^+^ cells in untreated samples and expressed in percentage. Data are shown as mean ± SEM of n>10 images, taken on 2 independent biological replicates. **e.** Schematic illustration showing the experimental procedure and timeline used to investigate the regenerative ability of NMO-rSkMs. **f**. Representative stereomicroscope images showing whole mount immunofluorescence staining for TUJ1 (red) and DESMIN (green) of NMO-rSkMs at day 35 of co-culture not treated with cardiotoxin (no CTX) or day 1, day 5 and day 20 after CTX treatment. Scale bar, 1 mm. **g**. Representative Z-stack confocal immunofluorescence images for TUJ1 (red) and DESMIN (green), showing a higher magnification of the corresponding stereomicroscope images. Scale bars, 200 μm. **h**. Representative Z-stack confocal immunofluorescence image of NMO-rSkM cross-sections stained for LAMININ (green) and embryonic MHC (magenta). Nuclei are counterstained with Hoechst (blue). Scale bars, 50 μm.

Based on these observations, and to verify the regenerative abilities of our assembloids, we injured 35-day NMO-rSkMs with cardiotoxin [24,39] (CTX) to damage mature muscle fibers and evaluate degeneration and self-regeneration after 1, 5 and 20 days from injury (Fig. 4e). Thus, we initially used morphometric analysis of whole mount immunostained samples to qualitatively estimate the muscular and neuronal compartment organization before damage, just after the injury and at longer time points. Already after 1 day from CTX treatment, it was possible to notice a decreased presence of elongated DES^+^ cells, especially in correspondence with the area closer to the scaffold, and a spotted and less extensive TUJ1^+^ neuronal axon network (Fig. 4f,g). Upon 5 days from CTX injury, we observed a slight amelioration in the muscular compartment, which presented thin elongated DES^+^ cells (Fig. 4f,g). Accordingly, the expression of eMHC confirmed the presence of newly formed myofibers after 5 days from the acute injury, indicating the beginning of muscle regeneration (Fig. 4h). On the contrary, the TUJ1^+^ neuronal axon network appeared still damaged 5 days after injury (Fig. 4f,g). However, this situation changed and improved as the days went by until a morphometric organization of the neuronal and muscular compartments closer to the one of an undamaged construct was reached 20 days after CTX administration (Fig. 4f,g).

### NMO-rSkM assembloids display functional muscle regeneration upon 20 days from acute damage

Following an acute injury, within a certain entity of damage, healthy SkM can spontaneously regenerate and rebuild functional tissue.[37,38] Based on the previous results, we further investigated the muscular and neural compartments of our functional NMO-rSkM assembloids before and after 1, 5 and 20 days from injury. As first, we confirmed and quantified the morphological changes observed upon injury and regeneration. In our samples, we verified that following CTX treatment, in NMO-rSkM assembloids the muscle compartment was clearly damaged after 1 and 5 days from the injury (0.63±0.03×10^6^ A.U. and 0.76±0.04×10^6^ A.U.), but restored to pre-CTX levels after 20 days as shown by the expression and quantification of DES^+^ signal (2.30±0.08×10^6^ A.U. and 2.12±0.10×10^6^ A.U.; Fig. 5a,b). When the neuronal compartment was investigated, we revealed that also this part was damaged upon CTX treatment, even though with delayed timing in respect to the muscular counterpart (Fig. 5c,d). As foreseen, a partial regeneration of neuronal protrusions and TUJ1^+^ neurofilaments occurs spontaneously but at a later time than muscle regeneration. Indeed, the mean axonal projection length decreased from 450.00±17.00 µm in no CTX samples to 35.06±2.97 µm 5 days after injury but started to increase by day 20 (274.12±13.25 µm) (Fig. 5c,d). Furthermore, the structural regenerative phase is anticipated by a prominent change in the proportion of proliferating PAX7^+^ stem cells which increase significantly in the first days after injury (24.20±2.70% on day 1 and 14.72±2.38% on day 5) returning to basal levels on day 20 (6.28±1.66% in the untreated sample and 5.46±1.40% on day 20; Fig. 5e). Importantly, the morphometric regeneration of the muscle compartment (Supplementary Fig. 6a) was confirmed at a functional level by the quantification of the contraction after ACh administration. Despite an evident decrease in the contractile capacity of NMO-rSKM assembloids after 1 and 5 days post-CTX injury (107.41±35.95 A.U. on day 1 and 278.41±57.34 A.U. on day 5), on day 20 the contraction levels returned to those measured before damage (1049.62±157.87 A.U. on day 20 vs 832.65±143.19 A.U. before damage; Fig. 5f, Supplementary Fig. 6b and Supplementary Video 4). According to the dynamics of neuronal and muscular recovery observed in damaged assembloids, we found rare juxtapositions of pre- and post-synaptic elements of the NMJ 20 days after injury (Fig. 5g). In agreement, we found that the injured NMO-rSkM could not reach a complete recovery of the NM system functionality after 20 days from CTX administration, as shown by partial ability of the assembloid to contract after administration of Glu (18.11±6.60 A.U. on day 1 vs 66.87±8.36 A.U. on day 5 and 165.66±24.81 A.U. on day 20; Fig. 5h, Supplementary Fig. 6c and Supplementary Video 5). These data confirmed that NMO-rSkM assembloids are able to respond to acute injury with SkM regeneration, and partial restoration of the NMJ functionality, suggesting a long-lasting integration between NMO and rSkM.

**Figure 5.**
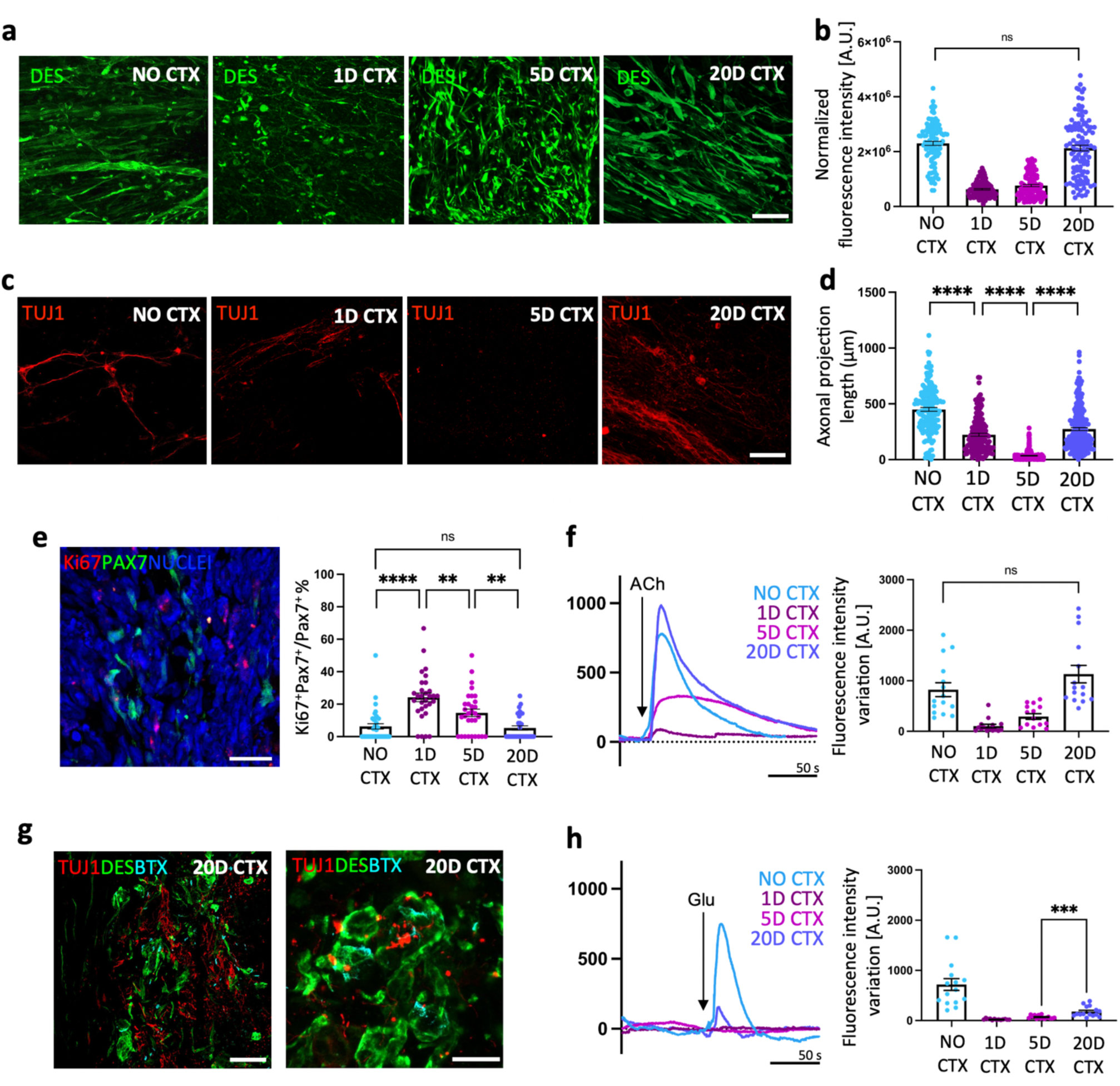
Functional muscle regeneration precedes neuronal and NMJ regeneration in NMO-rSkM assembloids. **a**. Representative Z-stack confocal immunofluorescence images of NMO-rSkMs at D35 of co-culture not treated with cardiotoxin (no CTX) or 1 day, 5 days and 20 days after CTX treatment and stained in whole mount for DESMIN (green). Scale bar, 50 µm. **b**. Quantification of total fluorescence intensity normalized for background signal performed on DESMIN immunofluorescence staining. Data are shown as mean ± SEM of 36 ROIs per biological sample per each condition (n=3). Statistical significance was determined using Mann-Whitney U test; ns, not statistically significant. **c**. Representative Z-stack confocal immunofluorescence images of NMO-rSkMs at day 35 of co-culture not treated with cardiotoxin (no CTX) or 1 day, 5 days and 20 days after CTX treatment and stained in whole mount for TUJ1 (red). Scale bar, 50 µm. **d**. Quantification of axonal projection length performed on TUJ1 immunofluorescence staining. Data are shown as mean ± SEM of >50 axonal projections per sample per each condition (n=3), measured on ≥ 3 images per condition. Statistical significance was determined using Mann-Whitney U test; ****P<0.0001. **e**. Left: Representative Z-stack confocal immunofluorescence image of NMO-rSkM at D35, stained in cryosections for PAX7 (green) and Ki67 (red). Nuclei are counterstained with DAPI (blue). Scale bar, 20 µm. Right: Quantification of Ki67^+^ cells among PAX7^+^ cells in NMO-rSkMs at day 35 of co-culture not treated with cardiotoxin (no CTX) or 1 day, 5 days and 20 days after CTX treatment, expressed in percentage. Data are shown as mean ± SEM of ≥ 30 images, from 2 different biological samples, per each condition. **f**. Left: Representative quantification of mean normalized fluorescence intensity variation registered during contraction of NMO-rSkMs at day 35 of co-culture not treated with cardiotoxin (no CTX) or 1 day, 5 days and 20 days after CTX treatment when stimulated with ACh. For each curve, data are shown as the mean of 3 independent replicates. Right: Contraction quantification of NMO-rSkM at D35 not treated with cardiotoxin (no CTX) or 1 day, 5 days and 20 days after CTX treatment. Contraction was recorded upon exogenous ACh administration, and is expressed as the maximum peak of fluorescence revealed via live imaging analysis. Quantifications were performed on 3 independent biological replicates per condition. Each of the 3 samples was divided into 5 ROIs and every dot represents a single ROI. Statistical significance was determined using Mann-Whitney U test; ns, not statistically significant. **g**. Representative Z-stack confocal immunofluorescence images of NMO-rSkMs cross sections 20 days after CTX treatment, stained for TUJ1 (red), DESMIN (green) and α-BTX^+^ regions (cyan). Scale bars, 50 µm (left) and 20 µm (right). **h**. Left: Representative quantification of mean normalized fluorescence intensity variation registered during contraction of NMO-rSkMs at day 35 of co-culture not treated with cardiotoxin (no CTX) or 1 day, 5 days and 20 days after CTX treatment when stimulated with Glu. For each curve, data are shown as the mean of 3 independent replicates. Right: Contraction quantification of NMO-rSkM at D35 not treated with cardiotoxin (no CTX) or 1 day, 5 days and 20 days after CTX treatment. Contraction was recorded upon exogenous Glu administration, and is expressed as the maximum peak of fluorescence revealed via live imaging analysis. Quantifications were performed on 3 independent biological replicates per condition. Each of the 3 samples was divided into 5 ROIs and every dot represents a single ROI. Statistical significance was determined using Mann-Whitney U test, ***P=0.0002.

## Discussion

In this work, we developed a novel in vitro platform to investigate and reveal the complex cellular and biological events involved during the integration of the human SkM and the neural system within an implantable tissue-engineered muscle construct by producing graft-host SkM assembloids.

Engineered SkM constructs hold the promise to be a therapeutic route for large and impactful muscle defects (single traumatic events, chronic conditions and exercise-induced injuries)[40,41] or congenital malformations (such as diaphragmatic hernia and abdominal wall defects).[42,43] However, their application in the clinic is still poor and it remains an issue of how to predict the efficiency of the developed engineered construct and establish the probability of success of engraftment and functional muscle regeneration in a humanized context. In addition, considering the application in the pediatric field for the treatment of congenital defects and malformations, it is extremely complicated to predict the response of a physiologically growing tissue/organ to the implantation of an engineered construct. Up to now, these aspects have been mainly investigated using various animal models.[4,44,45] Animals offer a valuable opportunity to gain insight into regenerative mechanisms and systemic responses to the graft; however, ethical limitations coupled with specie-specific unpredictable biological responses, restrict their application in predicting the behavior of host human cells in relation to the graft.[20,21]

Here we developed a humanized in vitro model where the muscular and neuronal contribution to the engineered SkM, and vice versa, could be studied in vitro. To do so, we combined hiPSC-based organoid technology and a well-established SkM tissue engineering approach to derive graft-host SkM assembloids. In the last few years, the production of NMOs enabled the long-term culture of a human NM system, allowing the investigation of interactions between MNs and SkM at the level of the NMJ in health and disease.[22–25,46] Therefore, NMOs are generally considered a promising approach for modeling in vitro human NM biology, diseases [23–25] and muscle regeneration.[24] Based on this, we hypothesized that NMOs could be used as an archetype of transplant acceptance or failure of human origin to replace animal experiments, especially for mimicking fetal/neonatal NM system that could mirror conditions of congenital malformations. We show that upon 22 days of hiPSCs differentiation, the derived NMOs presented the main NM system cell populations, from the expression of early and late myogenic markers to the presence of neuronal protrusion toward the internal muscular compartment. One of the most relevant aspects for the positive outcome of a tissue-engineered muscle graft is to understand whether the construct would integrate within the recipient tissue and vice versa, both from structural and functional points of view. In particular, in the SkM contest, the achievement of adequate and timely innervation by the host peripheral nervous system is mandatory to allow functional graft integration.[18] Here, we show that tissue-engineered products, such as dSkM and dSkM recellularized with hMPCs (i.e. rSkM), could be invaded, repopulated and remodeled by muscular and neuronal cell types derived from NMOs, generating graft-host SkM assembloids. These models allowed the in vitro evaluation of the interaction between the human NM system and engineered SkM constructs.

Until now, the most clinically relevant tissue engineering approach for volumetric SkM loss treatment has been performed using a decellularized urinary bladder scaffold.[15,17] Although treated patients recovered partial muscle volume and a certain degree of functionality, the study of Sicari and colleagues highlighted the importance of functional integration between the recipient tissue and the muscle graft for a positive outcome of the implant. We and other groups have demonstrated the importance of using tissue-specific scaffolds to obtain relevant results in repairing muscle damages and congenital defects, being the dSkM intrinsically equipped with muscle-specific and neurotrophic factors.[5,13,14] Our study confirms this concept: we found that the presence of a muscle-specific extracellular matrix allows cells to settle in anatomical positions more similar to those in vivo (e.g. under the myofiber basal lamina for satellite cells), increasing the degree of specificity of the regenerated SkM. In addition, transplantation of the single decellularized scaffold for large muscle defects is not a guarantee of complete and long-lasting treatment.[12,15,17] Accordingly, we found that although the dSkM alone is able to attract muscle and neuronal cells from the NMO, the host-cell invasion does not reach all areas of the scaffold in NMO-dSkM assembloids. On the contrary, the presence of pre-seeded hMPCs into dSkM (i.e. rSkMs) promotes a much more efficient and robust integration of cellular components from NMOs in NMO-rSkM assembloids. This was paralleled by significant improvement of the SkM contraction of the invaded tissue-engineered construct, strongly suggesting that repopulating dSkM with tissue-specific cells has a positive prediction to generate a functional graft once implanted in patients.

According to this, we also show that functional NMJs are generated upon the integration of the NMO-derived neural compartment with rSkMs in NMO-dSkM assembloids. Human NMJs have been shown to possess specific characteristics that distinguish them from comparable synapses in other mammalian species (e.g., mouse and rat).[47] These differences complicate the interpretation of animal studies for their applicability to humans. In our humanized in vitro 3D system, the presence of human NMJs formed by coin-shaped patches endplates [47] and capable of responding to neurotransmitters or toxins expectedly demonstrates their similarity to NMJs present in vivo, confirming the reliability of the model.

At the same time, the presence of stem and myogenic progenitor cells, such as PAX7^+^ cells identified in a satellite cell anatomical position, opened the possibility to investigate the long-lasting ability of the integrated tissue-engineered SkM construct upon damage. We used CTX, a well-established in vitro and in vivo assay to induce a transient and reproducible acute injury of the myofibers without directly affecting the nerves.[39] Our NMO-rSkM assembloids responded in a synchronous and coordinated manner to CTX, with degeneration and subsequent regeneration of the muscular compartment at first, and delayed degeneration and partial regeneration of the neuronal compartment. This experimental setup confirmed the effective integration between NMO and rSkM and that NMO-rSkM assembloids are able to undergo damage and respond with tissue regeneration in the expected times and ways.[48] Importantly, although SkM functionality was completely rescued after 20 days from the injury (i.e. upon ACh stimulation), the regeneration of the neuronal compartment with functional NMJs was still not fully achieved in the same timeline (i.e. upon Glu stimulation). External stimulation such as electrical, mechanical or magnetic forms, and other physical forces show promise in promoting in vivo and in vitro neural tissue regeneration.[49–51] After stimulation, neural tissue experiences increased proliferation, supporting axon sprouting and innervation. Thus, it could be of future interest to test whether external electrical stimulation could accelerate NM system regeneration in our assembloids. Despite the relevant results brought by our platform on the exploration of the interactions between the human NM system and engineered muscle tissue, it is important to underline that several key biological players are missing in the presented assembloid model. The future possibility to integrate into the NMO-rSkM assembloids other cell types, such as those belonging to the vascular tissue and the immune system, remains a fascinating perspective that can open new possibilities to dissect the regulation properties of other cellular players during the human NM system integration and regeneration.

In a broader scientific context in which researchers and clinicians increasingly seek to recreate 3D in vitro models able to mimic as much as possible the complex physiological environment, our NMO-rSkM assembloids can represent not only a preclinical test to predict the engraftment of specific muscle grafts but potentially also be a useful tool for studying drugs or therapeutic molecules for diseases that affect the NM system. The scientific community in the field of SkM and NM system research is strongly committed to the development of complex 3D in vitro models that can mimic myopathies and dystrophies,[52–56] or the effect of aging on physiological mechanisms of repair and regeneration.[57–59] In our 3D in vitro model, the presence of different cell types of the NM system, such as MNs, Schwann cells, mature muscle fibers and muscle stem cells will allow the in vitro acquisition of reliable results and facilitate the understanding of the events that occur in muscular and NM diseases and regeneration. Indeed, by implementing the NMO-rSkM model, we were able to mimic the human NM system degeneration/regeneration dynamics upon acute muscle injury, opening new perspectives to further investigate the relationship of the human NM system during degeneration and regeneration events.

Given its complexity in cellular composition, the high fidelity with the human context and the demonstrated functionality even after acute damage, our assembloids can open new studies on regeneration mechanisms both in healthy and diseased contexts, for a better understanding of translational applications in SkM-based regenerative medicine strategies.

## Acknowledgments

This work was supported by STARS Starting Grant 2017 of University of Padova (grant code LS3-19613), by Bando Direzione Scientifica IRP Città della Speranza (grant code 21/05) and by BIRD (code URCI_BIRD2223_01) to A.U.; by AFM Telethon (grant code) to N.E.; and by Bando CARIPARO-Ricerca Pediatrica 2020-2022 (grant number 20/17 FCR) to M.P. Schematic illustrations in Figures 1a, 2a, 2g, 3a, 4e were created with BioRender.com.

## Author contributions

A.U. and M.P. designed the study. L.R. performed the co-seeding experiments and analyses. L.R. and E.C. processed muscle biopsies, derived and characterized the hMPCs. L.R. and B.A. produced NMOs and performed molecular analysis. L.R. and M.L.P. performed the live imaging analyses. B.A., G.C. and P.C. contributed to sample preparation and imaging analysis. L.R. and E.M. derived dSkMs. O.G and S.A. produced and characterized the wild-type hiPSC line. C.La. produced and characterized the GFP hiPSCs. N.E. supervised the hiPSCs production and contributed to data interpretation. A.U. coordinated and supervised the NMO production and use. All the authors contributed to the revision of the manuscript. A.U. and M.P. analyzed and interpreted the data, wrote the manuscript and supervised the project.

